# Characterizing variability in resting-state functional magnetic resonance imaging (rsfMRI) metrics: a normative modeling framework

**DOI:** 10.64898/2026.05.28.728381

**Authors:** Alejandro Amador-Tejada, Ethan Danielli, Michael D. Noseworthy

## Abstract

Clinical adoption of new biomedical techniques depends on establishing reference values against which individual patients can be compared. In resting-state functional MRI (rsfMRI), most biomarker research has relied on the case-control paradigm, whose underlying assumptions are often invalid as diseases are frequently heterogeneous, limiting biomarker generalizability. Normative modeling offers a complementary alternative by characterizing individual deviations against a reference population. However, in rsfMRI, normative modeling has been applied almost exclusively to functional connectivity, with limited attention to age trajectories and sex effects. We address these gaps by developing a spatial normative model of four rsfMRI metrics that capture complementary features of the blood-oxygen-level-dependent (BOLD) signal across age and sex. Five publicly available datasets were aggregated to form a sample of 1,978 participants aged 10-30 years. Four metrics were computed for each of 110 grey matter regions: amplitude of low-frequency fluctuations (ALFF), fractional amplitude of low-frequency fluctuations (fALFF), regional homogeneity (ReHo), and Hurst exponent. A machine-learning model based on hierarchical Bayesian regression with a non-Gaussian likelihood was fitted per metric, modeling non-linear age effects, sex, and multi-site acquisition. Models were well calibrated across all four metrics, with fALFF showing the strongest predictive performance and Hurst exponent the weakest. Normative trajectories varied across brain regions for each metric, but on average, the median of each distribution remained bounded across regions, while the spread was more regionally variable. All four metrics showed predominantly negative slopes with age, indicating a decrease in each metric over the age window. This work provides a normative reference across four rsfMRI metrics that capture distinct features of the BOLD signal, complementing the case-control paradigm and supporting individual-level inference.

## Introduction

An important aspect of biomedical research is defining the clinical relevance of new techniques to encourage clinical adoption. This requires the establishment of expected reference baseline values to understand if a specific patient falls within a reference range. The case-control paradigm provides a method to investigate differences between the ‘average’ patient and a control group, providing group-level effect sizes as outcomes that are relatively simple to interpret and well understood across clinical and research environments [1]. This approach has become popular due to design and implementation simplicity, and cost-effectiveness [1]. However, this approach can inherently omit details, potentially hiding meaningful biological subtypes, individual patterns, or outliers within each group [2]. Furthermore, the case-control approach assumes that the groups being compared are internally homogeneous and distinguishable from one another [2,3]. Nevertheless, this assumption is often questionable as diseases are frequently heterogeneous [2,4,5], meaning two individuals can have the same diagnosis, yet different symptoms [4,6].

In response to the limitations, emerging approaches, such as normative modeling, a statistical framework to understand population heterogeneity, are now being considered. This framework defines a model that captures ‘normal’ variation in one or more biological or clinical measures within the population to establish a ‘normative’ frame of reference [2,7]. Normative modeling is a powerful tool because it can capture individual variations, allowing for the understanding of heterogeneity trends thereby enabling personalized assessments [7–9]. Furthermore, normative modeling represents a paradigm shift in disease conceptualization, framing disorders as significant deviations from typical patterns [2,7,9], which has been applied in psychiatry and cognitive neuroscience [8–12].

Resting-state functional MRI (rsfMRI) is a popular non-invasive MRI technique based on the blood-oxygen-level-dependent (BOLD) signal, used to study brain function at rest, in the absence of any task or stimulus [13,14]. It is a widely used technique due to its experimental simplicity: it does not require a study paradigm and involves minimal participant engagement, making it highly flexible [15,16]5/28/2026 8:37:13 AM. rsfMRI has become a valuable tool for identifying biomarkers of neurological and psychiatric disorders [17,18], which are increasingly recognized as having a biological basis, yet continue to be diagnosed exclusively through behavioral criteria [19,20]. Most research has relied mainly on functional connectivity (FC), regional homogeneity (ReHo), amplitude of low-frequency fluctuations (ALFF), and fractional ALFF (fALFF) [21]. Regardless of the rsfMRI-derived metric used, the case-control paradigm has remained the predominant framework for biomarker discovery across various diseases [18].

It is relevant to note that the case-control approach has several shortcomings, further reflected in the search for biomarkers using rsfMRI. For instance, most studies are characterized by relatively small sample sizes, heterogeneous populations, limited generalizability of findings, and proposed biomarkers with low disease specificity, as demonstrated by several meta-analyses [20,22–25]. Thus, while the case-control paradigm in rsfMRI offers value, it has faced challenges in producing clinically translatable biomarkers [5,18]. Normative modeling has been investigated as an alternative approach to address these challenges. However, only a limited number of studies have applied normative models to rsfMRI [19,26,27].

Beyond the scarcity of normative rsfMRI studies, large gaps in our understanding of brain health and function exist. A key gap is characterizing the healthy brain across the lifespan, as functional changes occur throughout life, with particularly pronounced shifts occurring during early development and in older age [28,29]. Furthermore, sex differences across the lifespan also affect many aspects of brain function that may or may not be regionally specific [30,31]. The omission of age and sex effects in the normative model design, an exclusive focus on FC analyses, and relatively small sample sizes persist in fMRI normative modeling research. Moreover, while FC is a widely used measure of brain function, it captures only a single feature of the BOLD signal. Numerous complementary features can also be extracted from the BOLD signal in rsfMRI [32]. Incorporating these diverse characteristics into normative modeling frameworks could support a more comprehensive and individualized characterization of deviations in brain function [27,33].

The field of brain rsfMRI research lacks a comprehensive understanding of normative modeling which includes sex and age, and other features of the BOLD signal [11,33], creating a research gap. Therefore, the main objective of this research was to develop a spatially normative model of rsfMRI that incorporates several features of the BOLD signal, enhancing the characterization of biological heterogeneity of brain function. Moreover, we argue that thoroughly characterizing normative populations is a critical prerequisite to studying disease. Establishing a reference framework could allow individual-level predictions, as normative modeling conceptualizes disease as a deviation from this normative baseline [5,19].

## Methods

### Data description

All data were pooled under local ethics approval, in addition to the ethical guidelines of each database and corresponding sites accessed. Five publicly available datasets (ABIDE-I [34], ABIDE-II [35], ADHD200 [36], FCP [37], and INDI [38]) were aggregated to create a larger rsfMRI sample. Detailed information for each database can be found in their respective publications. Beyond the specific criteria of each database, inclusion in the aggregated sample required participants aged 10-30 years, with available structural MRI and rsfMRI scans collected in 3T systems, and basic demographic data (age and sex). Participants aged 10-30 were chosen because this age range is common in neuroimaging research and in existing open-access datasets [39,40]. Furthermore, this age window has shown spatiotemporal changes in brain functional organization, with network-specific maturation peaking during adolescence and extending into early adulthood [41,42].

Participants with corrupted files or missing demographics were excluded from this study. To prevent participant duplication across databases, each dataset was cross-referenced against known repository overlaps documented in the literature, and a unified participant registry was maintained throughout data curation. For datasets with longitudinal or multiple acquisitions, only the baseline session was pooled.

### Data processing

rsfMRI preprocessing was performed with customized scripts using both <u>FSL</u> (v6.0.7.17) [43–45], and <u>AFNI</u> (v25.1.03) [46,47]. Preprocessing adhered to standard rsfMRI guidelines [21,32]. Briefly, analysis involved deletion of the first five functional volumes, slice timing and motion correction, spatial smoothing (Gaussian kernel, FWHM = 5mm), 4D global normalization, brain extraction, spatial normalization to the MNI152 T1 2mm template, and high-pass temporal filtering (0.01Hz). Quality assessment of preprocessed data included visual inspection of the raw data, co-registration, and assessment of spatial normalization accuracy. Subjects with excessive head motion (mean framewise displacement, FD > 0.5mm) were excluded from further analysis to minimize its impact on signal quality [48].

Following preprocessing, four rsfMRI metrics describing different features of the BOLD signal were computed for each subject: (1) amplitude of low-frequency fluctuations (ALFF) and (2) fractional ALFF (fALFF), depicting the frequency-based intensity of the BOLD signal; (3) regional homogeneity (ReHo), indicative of local connectivity; and (4) Hurst exponent, reflecting BOLD signal autocorrelation. Specific additional preprocessing steps were applied to each metric as required [16,32]. ALFF, fALFF, and ReHo maps were retained in their raw units, without applying typical within-subject normalization. After processing, each rsfMRI parametric map was parcellated into 110 grey matter regions of interest (ROIs) using the Harvard-Oxford cortical and subcortical atlases, and the mean value within each ROI was extracted and used as the input feature for normative modeling.

### Data analysis

The Predictive Clinical Neuroscience Toolkit (PCNToolkit) (version 1.1.2, Python 3.11), an open-source machine-learning package for normative modeling in neuroimaging and psychiatry [33,49], was used to model the normative regional variation of these rsfMRI metrics.

Briefly, the PCNToolkit uses a Bayesian regression framework. For each rsfMRI response variable (y), the model takes the form:

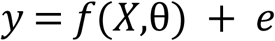

Where X are the covariates (e.g., age, sex), θ are model parameters and e is the residual. Bayesian inference combines prior beliefs 𝑝(𝜃) with the likelihood 𝑝(𝑦 | 𝑋,𝜃) via Bayes’ theorem to yield the posterior distribution: 𝑝(𝜃 | 𝑋,𝑦) ∝ 𝑝(𝑦 | 𝑋,𝜃) 𝑝(𝜃).

A Hierarchical Bayesian Regression (HBR) model with Sinh-ArcSinh (SHASHb) likelihood function (i.e., HBR-SHASHb) was used with a B-spline basis expansion (degree=3, knots=3) to model non-linear effects of age. The HBR model accounts for the multi-site nature of the aggregated rsfMRI sample via partial pooling in which site-specific parameters (θ_i_) are drawn from shared distributions across sites. Random intercepts were modelled per site for both the location (μ) and scale (σ) to account for scanner effects [50,51]. Moreover, the SHASHb likelihood, 𝑝(𝑦 | 𝑋,𝜃) = 𝑆𝐻𝐴𝑆𝐻𝑏 (𝜇(𝑋), 𝜎(𝑋), 𝜀, 𝛿), extends the standard Gaussian HBR model by modeling skewness (ε) and tail weight (δ) alongside μ and σ, allowing for flexible non-Gaussian predictive distributions [52].

Weakly informative priors were specified for each rsfMRI metric in the absence of established prior distributions, guided by the theoretical properties and small-sample empirical distributions of each metric [32,37,53]. A detailed description of the PCNToolkit, its implementation, model setup, evaluation and deviation scores computation is openly available [33,50,52,54].

Normative models were built for each rsfMRI of the four metrics. Each model had 110 response variables corresponding to the rsfMRI metric average value for each ROI, with two covariates (age, sex) and one batch effect (site) to also account for differences in rsfMRI sequence parameters. An 80/20 train/test split, with stratification based on batch effects, was employed. Five evaluation statistics were used to assess the normative models on the validation set: (1) explained variance (EXPV); (2) Pearson’s correlation coefficient (RHO); (3) Standardized mean squared error (SMSE); (4) Shapiro-Wilk test; and (5) Mean standardized log loss (MSLL) [33,50,55].

Normative trajectories were visualized by generating predictions on a synthetic covariate grid spanning the age window, crossed with sex and batch effects. Centiles were averaged across sites to produce a population-level normative curve for each brain region. For illustrative purposes, trajectories are shown for four brain regions: Caudate (Right, CAU), Frontal Pole (Left, FP), Insular Cortex (Right, INS), and Lateral Occipital Cortex inferior division (Right, iLOC). These regions were selected because they encompass the peak MNI coordinates of brain regions that showed significant transdiagnostic alterations in clinical populations compared to controls, as reported in an rsfMRI meta-analysis of studies using ALFF, fALFF, and ReHo [56]. The normative trajectories for all brain regions are provided in the supporting material.

Descriptive statistics were used to examine central tendency and dispersion stratified by sex, and the effects of age on normative trajectories. Lastly, a visual representation of the spatial distribution of these metrics was generated by averaging across the age window and sex. The boundaries of the default mode network (DMN) nodes, extracted from the CONN toolbox [57,58], were overlaid to facilitate visualization of the spatial correspondence between the DMN regions and the observed metric distributions.

## Results

Subjects meeting one or more exclusion criteria (n=67), exceeding the FD threshold of 0.5 mm (n=61), and failing processing quality assessment due to co-registration issues (n=52) were excluded from further analysis. A total of 1,978 participants between the ages of 10-30 years old were included in this study. **Fig 1** shows the number of participants and the proportions of males and females across the age window (**Fig 1a**) and for each site (**Fig 1b**). In addition, **S1 Table** describes in detail the contributing databases and sites from which participants were drawn to form the aggregated rsfMRI sample. For each of the five databases and their corresponding sites, this table summarizes the sample size, sex distribution, MRI manufacturer, and rsfMRI sequence parameters, including TR, TE, number of time points, voxel size, number of slices, and eye condition.

**Figure 1:**
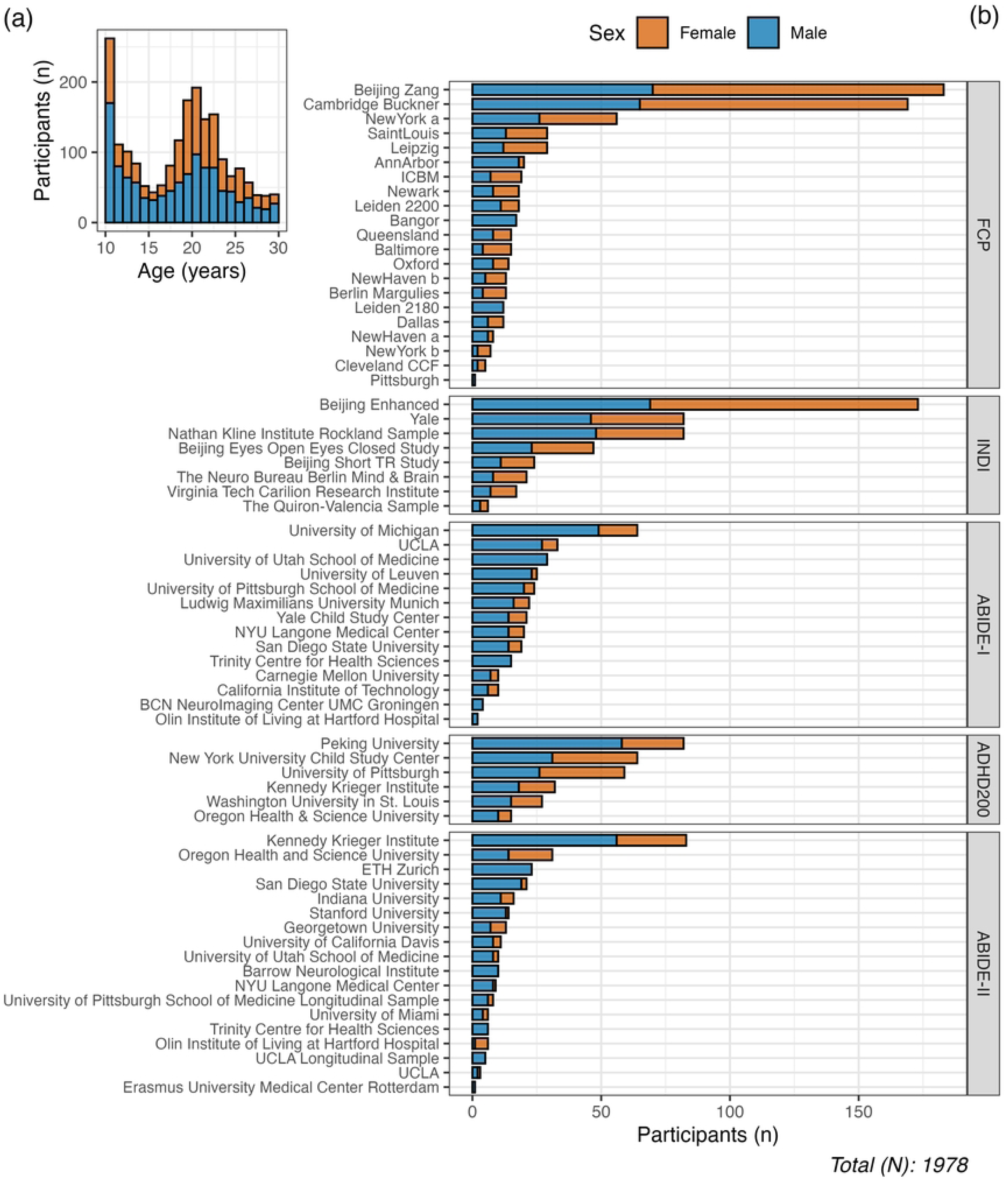
Summary of the number of participants and the proportion of males (blue) and females (orange) pooled (a) across the age window and (b) from each site within each database.

Four rsfMRI metrics were computed and averaged for each of the 110 ROIs, yielding 440 observations per participant. Next, four normative models were built, one per rsfMRI metric: ALFF, fALFF, ReHo, and Hurst exponent. Each model used age and sex to predict the rsfMRI metric value for each ROI, accounting for site variability, employing an HBR model with a SHASH likelihood.

The evaluation statistics from the validation set are shown in **Fig 2**, with a more detailed summary shown in **S2 Table**, and the interpretation and ideal values of each statistic are shown in **Table 1**. On average, fALFF showed higher EXPV and RHO than ALFF and ReHo, while Hurst exponent showed the lowest values across both statistics. Conversely, fALFF showed lower SMSE and MSLL, followed by ALFF and ReHo, while Hurst exponent showed the highest values for both statistics. For all four metrics, ShapiroW statistic values were close to 1.

**Figure 2:**
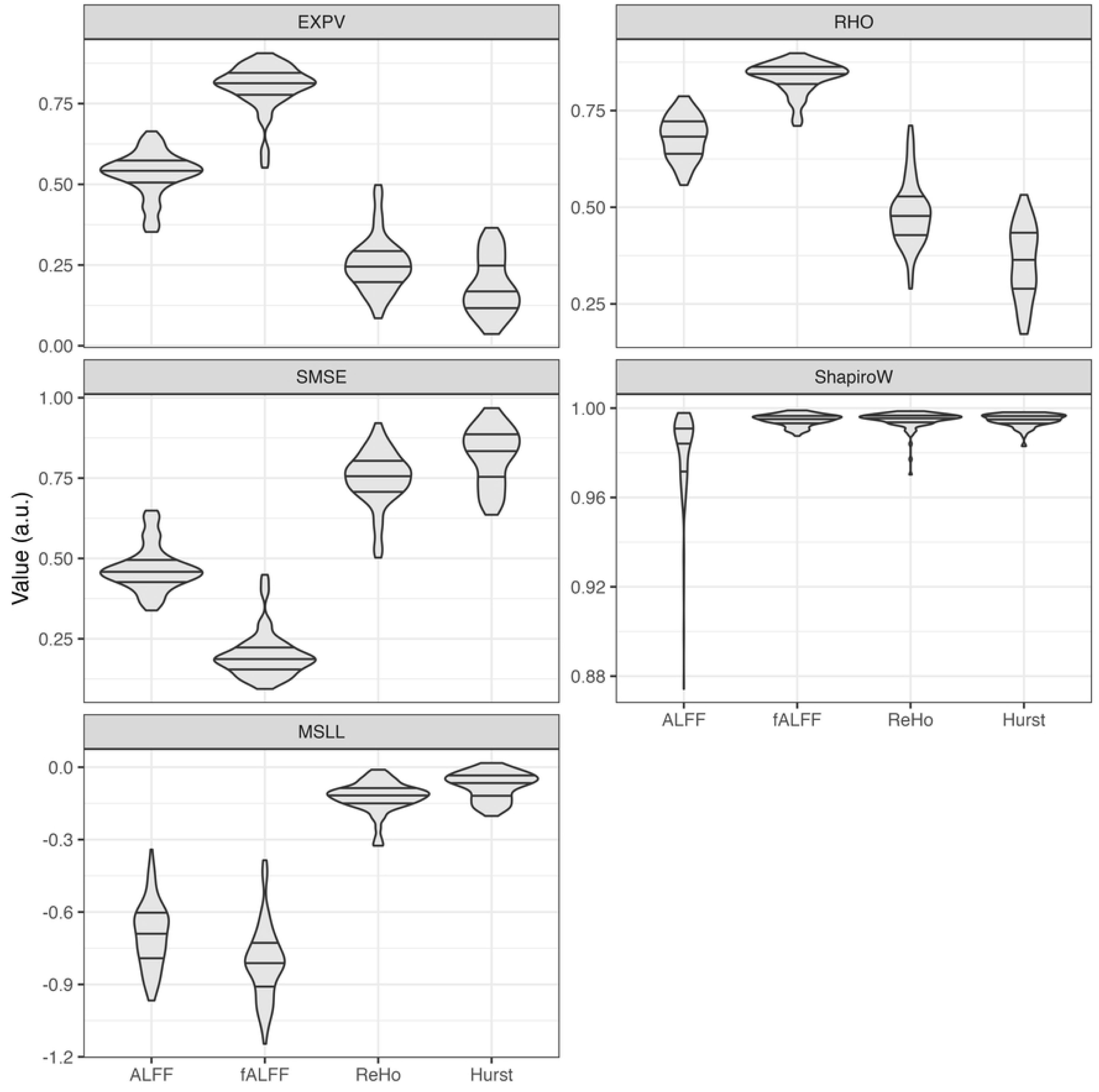
Evaluation statistics for each rsfMRI metric normative model. The reported statistics are (1) explained variance (EXPV); (2) Pearson’s correlation coefficient (RHO); (3) Standardized mean squared error (SMSE); (4) Shapiro-Wilk (ShapiroW) test; and (5) Mean standardized log loss (MSLL). The interpretation of these statistics is shown in **Table 1**.

**Table 1:**
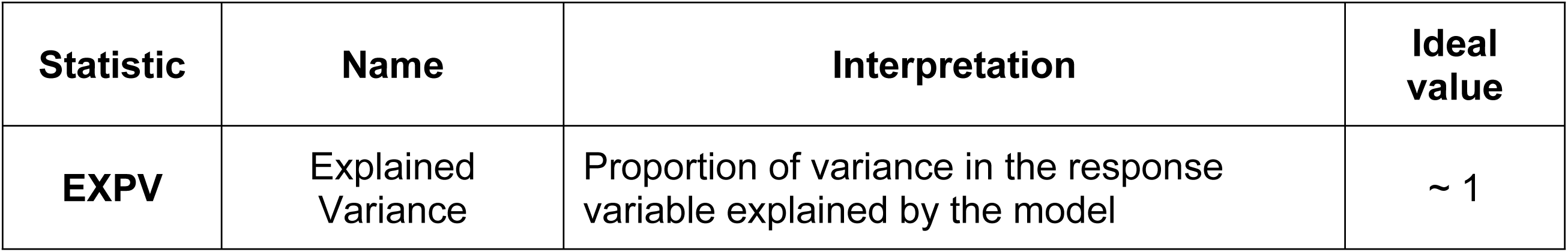
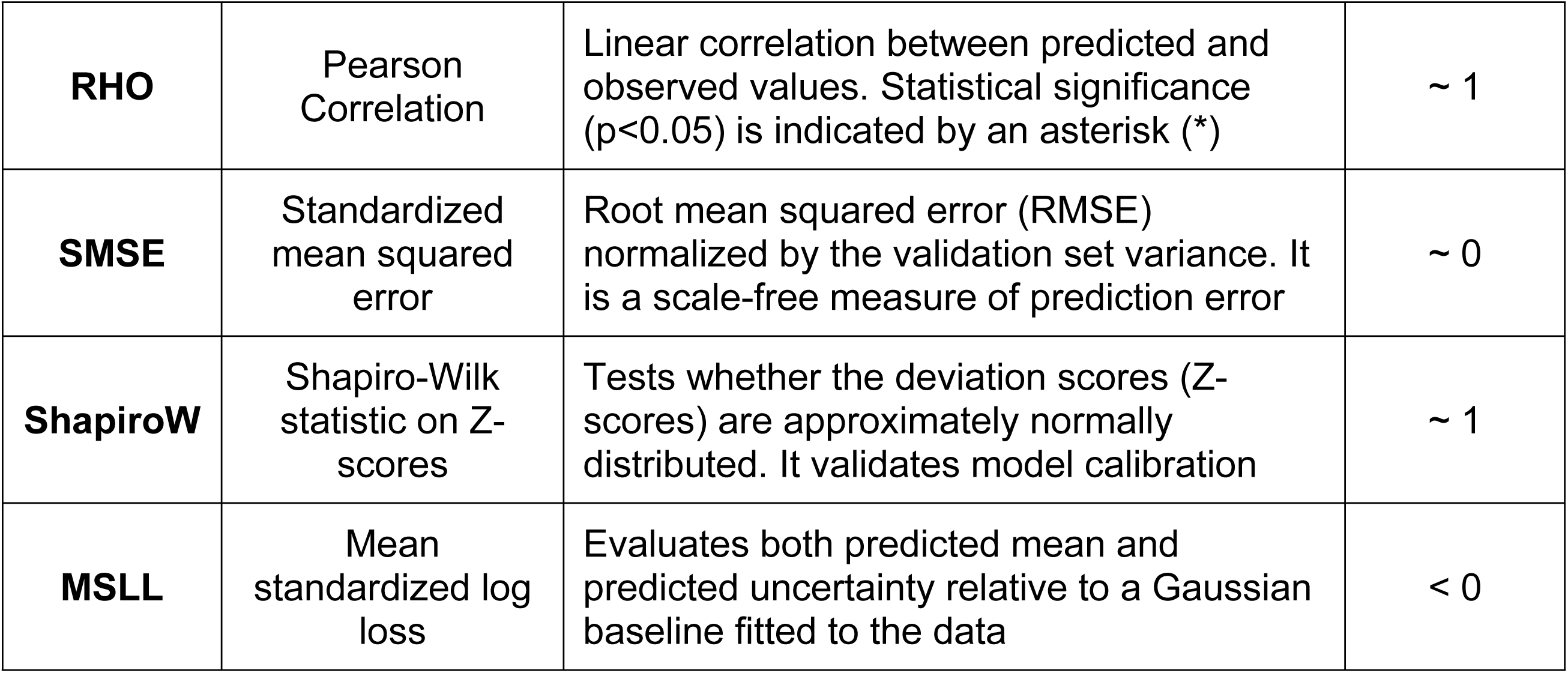
Summary of the five evaluation statistics reported in this study, including their interpretations and ideal values [33,50].

The trajectories for the four rsfMRI metrics, across the four selected ROIs, are shown in **Fig 3**. Each plot shows the 5^th^, 25^th^, 50^th^ (median), 75^th^, and 95^th^ centile curves against age, coloured by sex. Similarly, the trajectories for all brain regions are shown in **S1 Fig**. We observed that each metric has a distinct normative trajectory across age, varying between regions.

**Figure 3:**
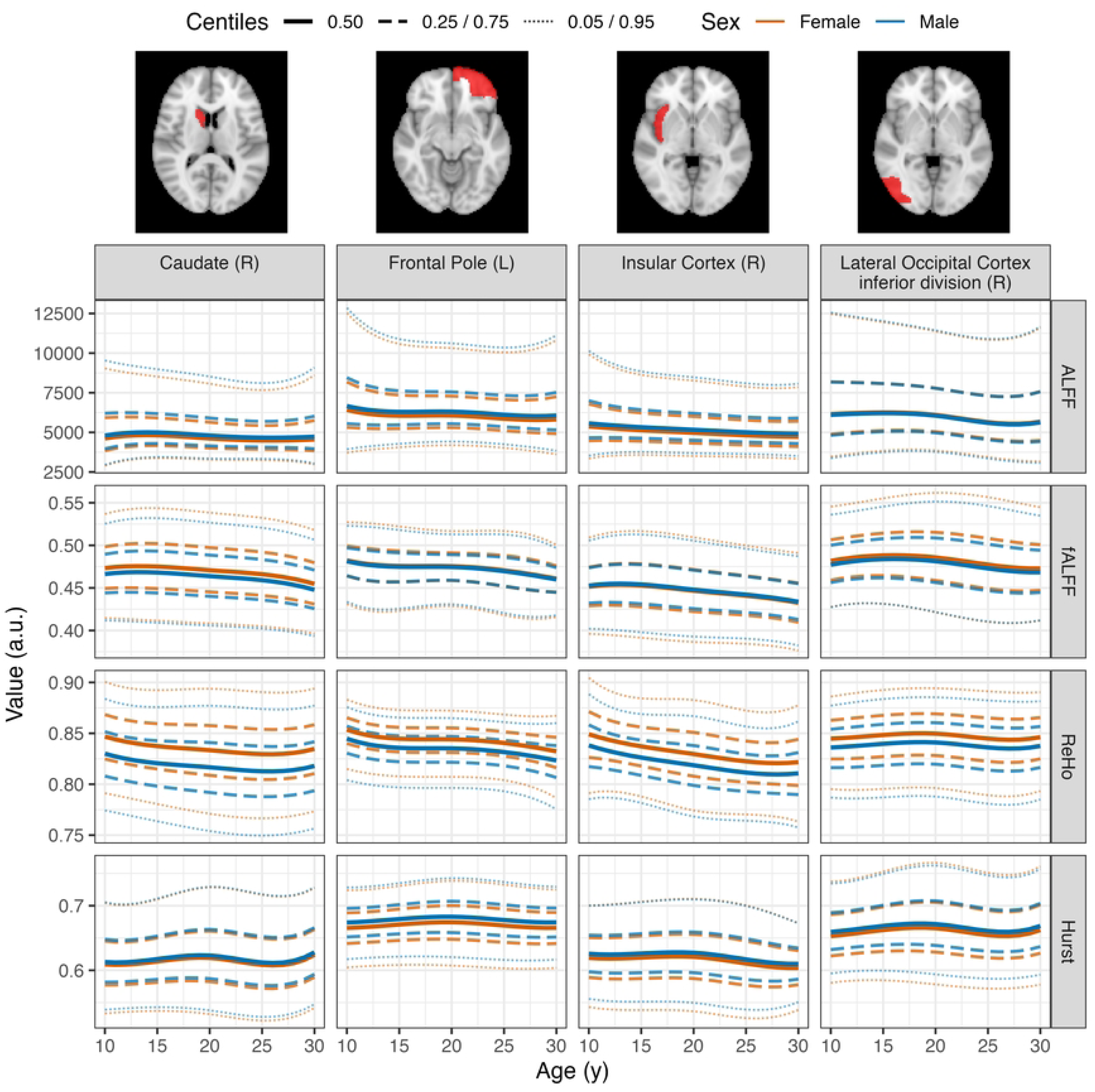
Normative trajectories across age for each rsfMRI metric (rows) for four ROIs (columns). Five centile curves are shown: 5^th^ and 95^th^ (dotted line, thin line width); 25^th^ and 75^th^ (dashed, medium line width), and 50^th^ (i.e., median; solid, thick line width). Sex-specific centile curves are displayed, with blue denoting males and orange denoting females. The normative trajectories for all brain regions across the four metrics are shown in **S1 Fig**. L: left; R: right.

**Fig 4** characterizes the center, spread, and age effects of the normative trajectories for all ROIs. **Fig 4a** presents the distribution of the median (p50) averaged across age, while the age-averaged 90% reference interval (i.e., p95 – p05) is seen in **Fig 4b**, both stratified by sex. For both sexes, all metrics showed median variation across brain regions: ALFF had the widest interquartile range (IQR), while fALFF, ReHo, and Hurst had narrower IQRs. No outliers were observed using a threshold of 1.5 IQR above or below the distribution of medians (**Fig 4a**). The 90% reference interval also varied across regions, with wider intervals on average in females than males for fALFF, ReHo, and Hurst exponent. But, in contrast to the median, outliers were identified (**Fig 4b**). ALFF showed the highest number of ROIs whose reference interval exceeded 1.5 IQR above the distribution of all intervals, followed by fALFF, and then ReHo. In contrast, Hurst exponent showed narrow reference intervals for all brain regions. Regions whose interval widths exceeded 1.5 IQR were labeled as outliers and are shown in **Table 2**. **Fig 4c** displays the slope of linear fits applied to the 50^th^ centile (i.e., the median, p50) as a function of age. All brain regions in ALFF showed negative slopes, while the average slope is negative for fALFF, ReHo, and Hurst exponent.

**Figure 4:**
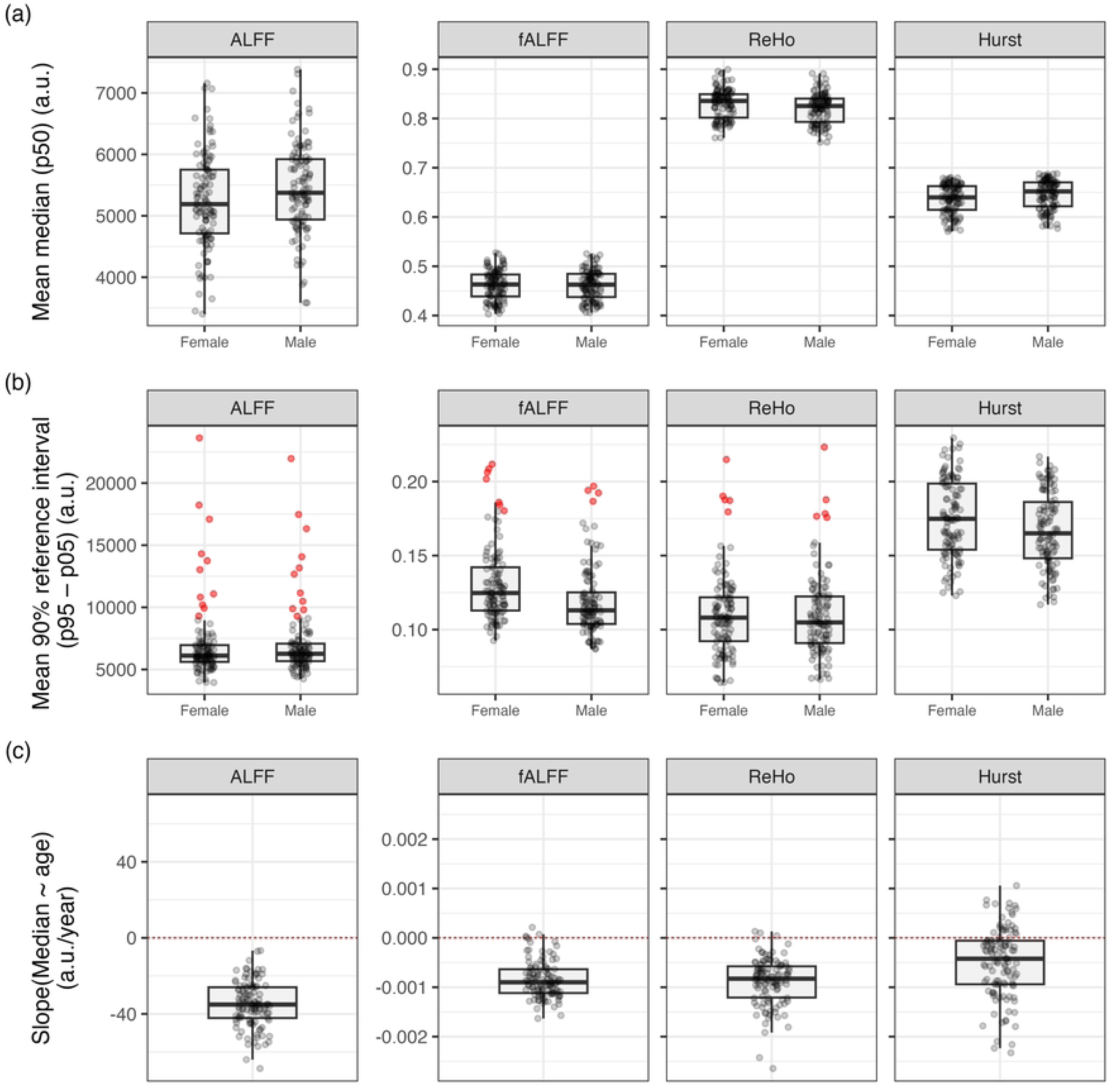
Summary statistics of normative trajectories for the four rsfMRI metrics. Distribution of the (a) median (p50) and (b) 90% reference interval (p95 – p05) for all ROIs, averaged across age, split by sex. Regions whose reference interval exceeded 1.5 IQR above the distribution of all intervals are shown in red (a,b). (c) Distribution of the slopes from linear fits on the 50^th^ centile (i.e., the median, p50) as a function of age for all ROIs. A red dashed line is shown in (c) to differentiate positive and negative slopes. For visualization purposes, fALFF, ReHo, and Hurst exponent share a common y-axis scale to facilitate direct comparison, while ALFF is displayed on an independent scale due to its characteristics.

**Table 2:**
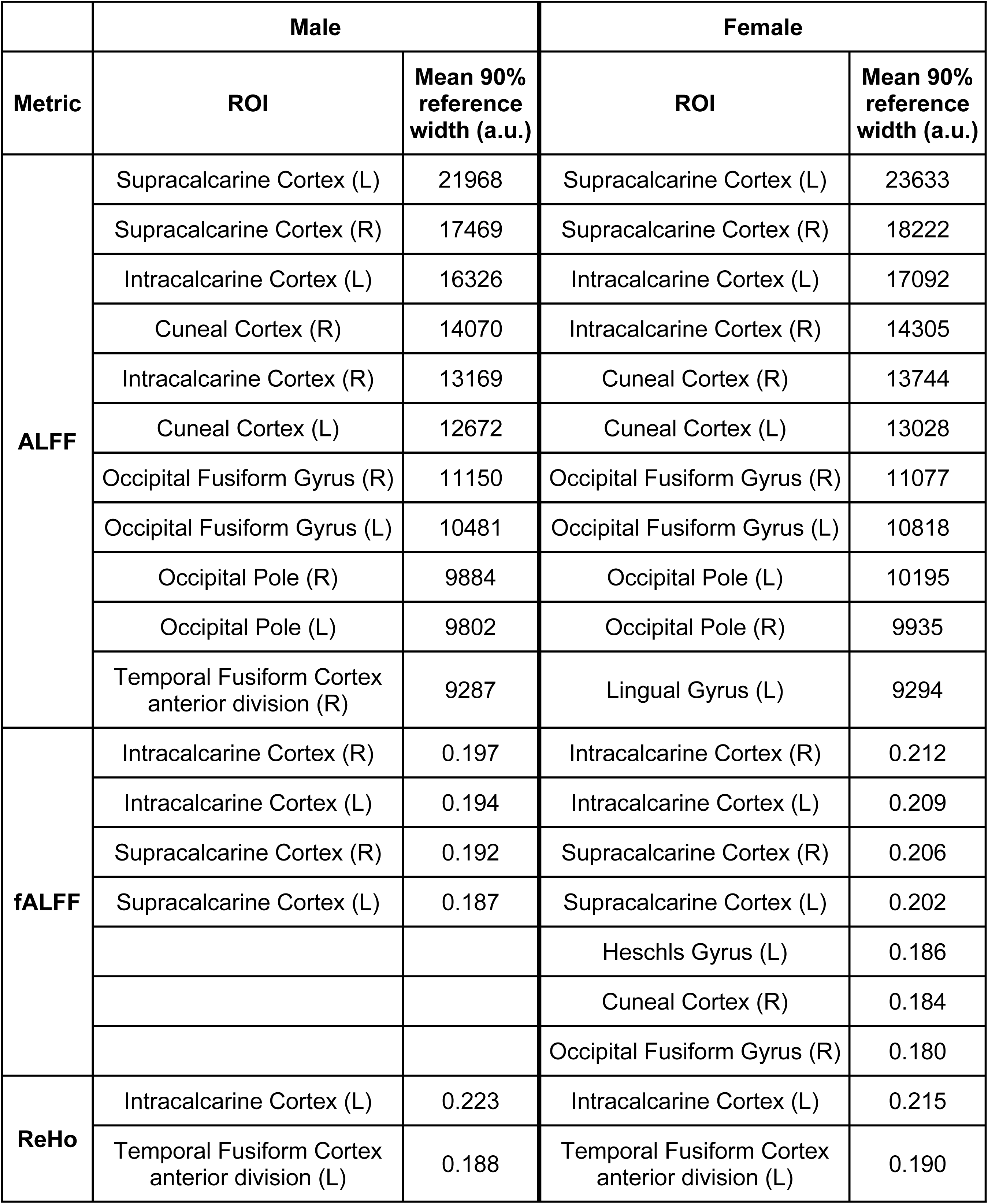
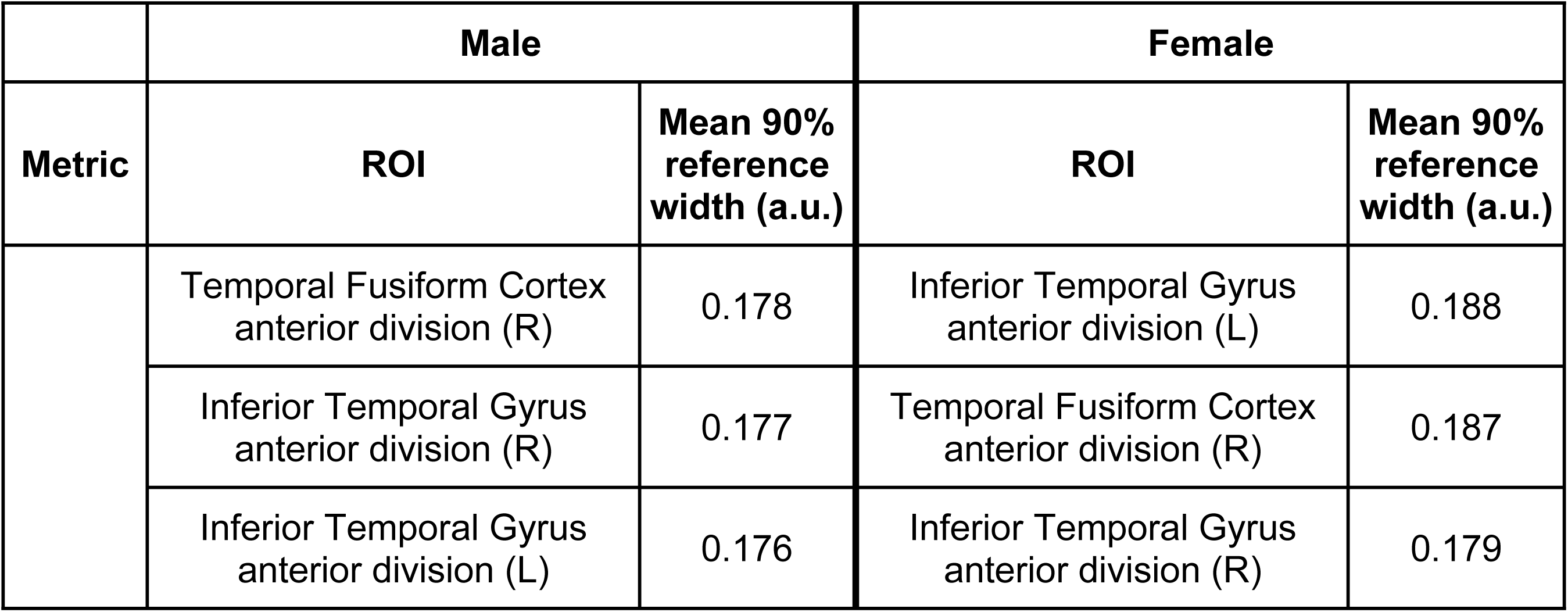
List of ROIs whose 90% reference interval (p95 – p05) was labelled as outliers by having a width interval higher than 1.5 IQR above the distribution of all intervals. Of note, Hurst exponent showed narrow reference intervals for all brain regions and hence no outliers were identified. L: left; R: right.

|  | Male |  | Female |  |
| --- | --- | --- | --- | --- |
| Metric | ROI | Mean 90%<br>reference<br>width (a.u.) | ROI | Mean 90%<br>reference<br>width (a.u.) |
| ALFF | Supracalcarine Cortex (L) | 21968 | Supracalcarine Cortex (L) | 23633 |
|  | Supracalcarine Cortex (R) | 17469 | Supracalcarine Cortex (R) | 18222 |
|  | Intracalcarine Cortex (L) | 16326 | Intracalcarine Cortex (L) | 17092 |
|  | Cuneal Cortex (R) | 14070 | Intracalcarine Cortex (R) | 14305 |
|  | Intracalcarine Cortex (R) | 13169 | Cuneal Cortex (R) | 13744 |
|  | Cuneal Cortex (L) | 12672 | Cuneal Cortex (L) | 13028 |
|  | Occipital Fusiform Gyrus (R) | 11150 | Occipital Fusiform Gyrus (R) | 11077 |
|  | Occipital Fusiform Gyrus (L) | 10481 | Occipital Fusiform Gyrus (L) | 10818 |
|  | Occipital Pole (R) | 9884 | Occipital Pole (L) | 10195 |
|  | Occipital Pole (L) | 9802 | Occipital Pole (R) | 9935 |
|  | Temporal Fusiform Cortex<br>anterior division (R) | 9287 | Lingual Gyrus (L) | 9294 |
| fALFF | Intracalcarine Cortex (R) | 0.197 | Intracalcarine Cortex (R) | 0.212 |
|  | Intracalcarine Cortex (L) | 0.194 | Intracalcarine Cortex (L) | 0.209 |
|  | Supracalcarine Cortex (R) | 0.192 | Supracalcarine Cortex (R) | 0.206 |
|  | Supracalcarine Cortex (L) | 0.187 | Supracalcarine Cortex (L) | 0.202 |
|  |  |  | Heschls Gyrus (L) | 0.186 |
|  |  |  | Cuneal Cortex (R) | 0.184 |
|  |  |  | Occipital Fusiform Gyrus (R) | 0.180 |
| ReHo | Intracalcarine Cortex (L) | 0.223 | Intracalcarine Cortex (L) | 0.215 |
|  | Temporal Fusiform Cortex<br>anterior division (L) | 0.188 | Temporal Fusiform Cortex<br>anterior division (L) | 0.190 |
| Metric | ROI | Mean 90% reference width (a.u.) | ROI | Mean 90% reference width (a.u.) |
|  | Temporal Fusiform Cortex anterior division (R) | 0.178 | Inferior Temporal Gyrus anterior division (L) | 0.188 |
|  | Inferior Temporal Gyrus anterior division (R) | 0.177 | Temporal Fusiform Cortex anterior division (R) | 0.187 |
|  | Inferior Temporal Gyrus anterior division (L) | 0.176 | Inferior Temporal Gyrus anterior division (R) | 0.179 |

Lastly, **Fig 5** shows the spatial normative distribution of the median for each rsfMRI metric, averaged across age and sex. The age- and sex-averaged whole-brain mean, and IQR for each rsfMRI metric are also shown in **Fig 5**, with fALFF, ReHo, and Hurst exponent exhibiting narrower IQRs than ALFF.

**Figure 5:**
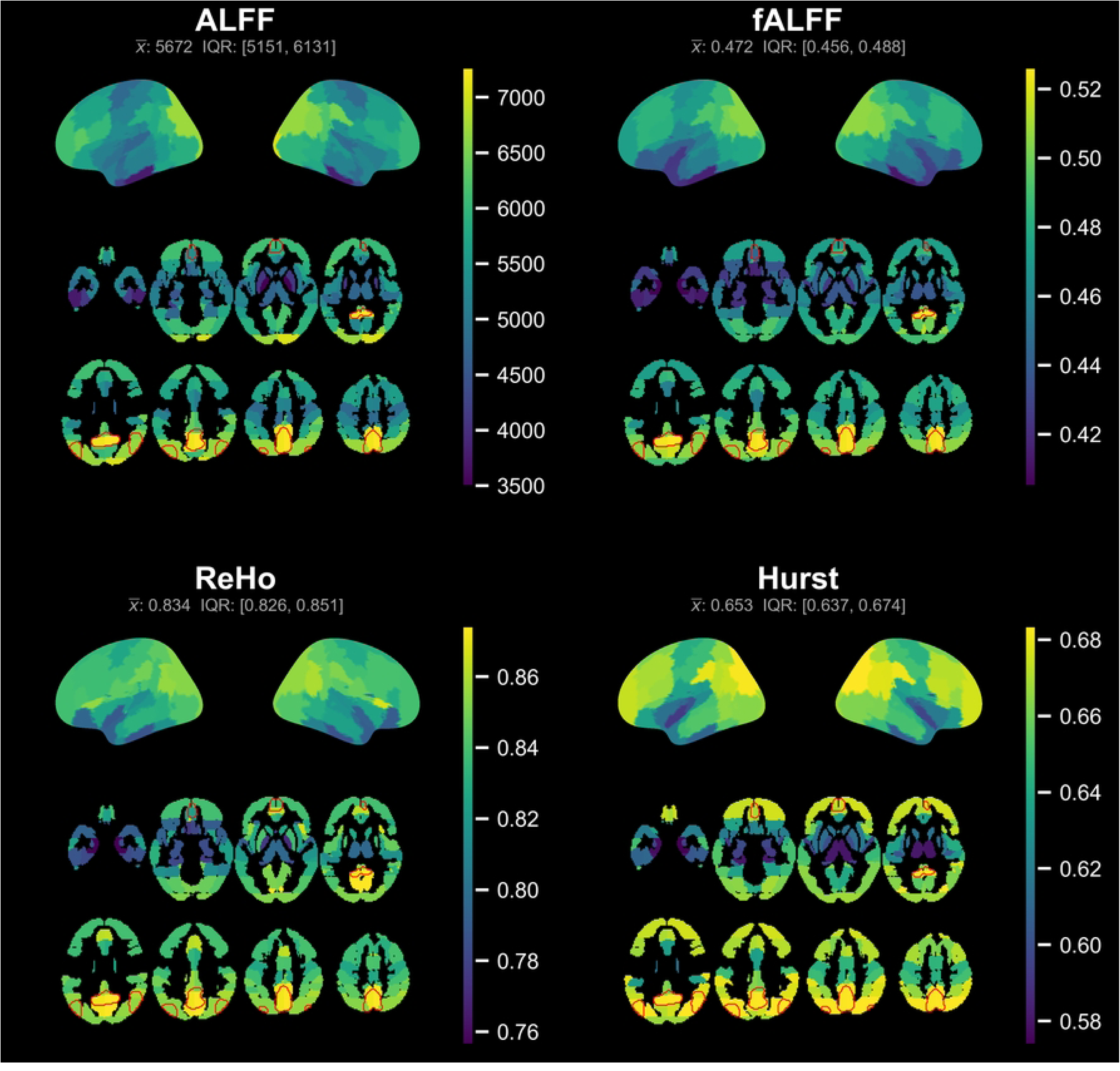
Spatial normative map of the age-averaged the 50^th^ centile (p50) for each rsfMRI metric, alongside the corresponding global mean and IQR, displayed on cortical surface projections and axial slices. The boundaries of the DMN nodes are shown in the axial montage, in red.

## Discussion

This research estimated spatial normative trajectories of four common rsfMRI metrics (ALFF, fALFF, ReHo, and Hurst exponent) that reflect age- and sex-related variation across brain regions, while accounting for MRI scan-related effects. Sex was included as a linear coefficient on the mean, without interacting with the age B-spline. This is consistent with prior sex-stratified analyses indicating that male and female normative trajectories share a common shape and differ only by a vertical offset [59].

The HBR-SHASH model performed differently across metrics, as shown in **Fig 2**. EXPV, RHO, and SMSE showed moderate-to-good performance, reflecting the ability of the model to predict the response variable with low-to-moderate error. The moderate performance of ReHo and Hurst exponent may reflect limited variance explained by age and sex covariates [51], a result also observed in normative modeling of FC [9]. However, ShapiroW and MSLL showed good performance across all metrics, indicating that the models are well-calibrated (Z-scores are approximately normal) and yield better predictions than a Gaussian model [33,50,55]. Although, five brain regions showed worse predictions than a Gaussian baseline (i.e., MSLL > 0) for Hurst exponent (Inferior Temporal Gyrus anterior division (L), Parahippocampal Gyrus posterior division (L), Heschls Gyrus (R), Occipital Fusiform Gyrus (R), Planum Polare (R)), suggesting that the predictive variance in these regions may be poorly calibrated [51,60].

The normative trajectories differed across ROIs as a function of age and sex, as seen in **S1 Fig** and **Fig 3**. These results are consistent with previous research that demonstrated spatial variations in the resting-state BOLD signal [13,37], indirectly reflecting the functional topography of the brain due to the neurovascular coupling [14,61]. Furthermore, previous research showed that ALFF, fALFF, and ReHo exhibit spatial patterns related to physiology, as evidenced by their correspondence with PET-derived measures of cerebral metabolic rate and hemodynamics [62]. The normative trajectories show modest differences between females and males, consistent with prior reports of sex-related variation in rsfMRI [37,63], typically attributed to physiological differences [64] (further summarized in **Fig 4a**).

Despite these spatial differences, the age-averaged median trajectory across ROIs showed no outliers across metrics, indicating the center of the distribution is relatively bounded across brain regions (**Fig 4a**). However, the 90% reference interval showed outliers for ALFF, fALFF, and ReHo (**Fig 4b**), meaning there are brain regions that have a wider distribution. This could be attributed to the susceptibility of these metrics to physiological noise, inter-subject biological variability, or model uncertainty [37,65].

The slope from a linear regression of the median trajectory indicates the overall direction of change across the age window. The slopes for all brain regions for the four metrics are shown in **Fig 4c**. The overall slope trend is negative, although the proportion of ROIs showing negative and positive slopes varies for each metric. This trend indicates that all metrics decrease with age within the time window. As sex was included as a linear coefficient on the mean, the age slope of the median is equivalent across sexes by construction and was not visualized separately in **Fig 4c**. Previous research has found widespread decreases in ALFF, fALFF, and ReHo with increasing age [66–69], which has been attributed to increasing functional integration and network specialization as part of maturation [66,68,69]. Similarly, Hurst exponent showed a negative trend consistent with previous research [70]. A lower Hurst exponent indicates higher BOLD signal complexity, which is associated with a greater capacity of the brain to transition between functional states [71].

Each rsfMRI metric measures a distinct characteristic of the BOLD signal [15,21,32]: ALFF and fALFF reflect frequency-based intensity, ReHo displays its local connectivity, and Hurst exponent shows its complexity. This is further reflected in the different mathematical formulations used to derive each metric [32]. Thus, it is expected that each metric will show a particular range of values as shown in **Fig 5**. The GM ranges reported in this research are within theoretical ranges for ReHo and Hurst exponent, reporting GM values within 0.6-0.8 for ReHo [72] and 0.5-0.8 for Hurst exponent [70,73]. Nonetheless, ALFF and fALFF are challenging to compare with previous research because they are typically reported as normalized values for group-level analyses [37,65]. To the authors’ knowledge, no normative raw reference ranges have been reported in the literature. Although, previously reported spatial patterns were observed, such as high ALFF and fALFF values in the DMN nodes [65].

### Limitations

The HBR-SHASH model used in this research is a robust approach that allows to (1) model non-Gaussian distributions [52,60]; (2) estimate predictive uncertainty [2]; (3) perform multi-site modeling [50]; and (4) use federated (i.e., decentralized) model learning [33,51]. Nevertheless, this approach has some disadvantages worth mentioning.

First, the models are estimated independently for each response variable (i.e., per brain region), so inter-regional correlations are ignored [51]. For instance, these normative models could omit local spatial patterns in ReHo, as it reflects local BOLD synchrony between a voxel and its neighbors. Moreover, HBR requires prior distributions over model parameters as part of the Bayesian inference pipeline [53]. Normative modeling has been predominantly applied to structural neuroimaging [59,74–77], and as such, there are no established reference parameters for rsfMRI metrics, hindering the use of strongly informative priors. Therefore, we used weakly informative priors guided by a preliminary characterization of rsfMRI metrics in a small dataset [32]. This approach constrains the posterior distribution from improbable parameter values [53] while still being flexible [51]. Nonetheless, the model could potentially underperform compared to strongly informative priors, particularly for sites with small sample sizes.

Another limitation is related to using a B-spline basis function. A relatively narrow age window was used to build normative trajectories of rsfMRI metrics, using a B-spline basis expansion with 3 degrees and 3 knots to model non-linear effects of age on these metrics. The choice of degrees and knots in a B-spline basis is inherently arbitrary, with no consensus on their specification. This issue can be understood with the bias-variance trade-off: too few knots or low polynomial degrees risk underfitting the data, producing a model with high bias but low variance that fails to capture non-linear behaviours. Contrarily, too many knots or high degrees risk overfitting, yielding models with low bias but high variance that are more sensitive to noise than signal [78,79]. Moreover, a single B-spline configuration applied uniformly across brain regions may be inappropriate. Previous research has shown intra-regional differences in the developmental trajectories of FC across age, showing different patterns: U-shaped, inverted U-shaped, and positive or negative linear trends [80,81]. A comprehensive investigation on the effects of age and model flexibility is needed to properly understand the linearity or non-linearity of rsfMRI metrics across brain regions.

An additional key limitation is the definition and setup of batch effects. A batch effect is typically understood as a category that introduces unwanted non-biological variation to the data. In neuroimaging, this variation is typically associated with the scanner characteristics and acquisition parameters [82]. This would require including several categories as batch effects, such as site, scanner manufacturer and model, and acquisition parameters (e.g., TR, TE, time points, etc.). Nonetheless, there is an important trade-off to consider: increasing the number of batch categories reduces the sample size per batch combination, which has been shown to decrease model stability and calibration accuracy [83]. In practice, multi-site neuroimaging studies treat the site as the main batch effect, as each site typically has unique and consistent hardware and acquisition parameters [59,75,76,84]. A further limitation arises with the use of open-access databases, where documentation, such as acquisition parameters, is inconsistent (see **S1 Table**). This impedes the use of more categories as batch effects, potentially leaving residual non-biological variance in the data. In such cases, we believe the most viable option is the use of the site as the main batch effect, yet it is important to recognize this is a limitation, as rsfMRI metrics have been shown to be sensitive to acquisition parameters [85,86]. Further work with larger pooled samples and more accurate documentation is needed to increase the number of batch effects and removal of non-biological variance, while increasing the number of subjects per batch combination.

## Conclusion

In summary, this work demonstrates the feasibility of building normative models in rsfMRI, establishing a spatial normative framework that allows individual-level statistical inferences, complementing, rather than replacing, the already well-established case-control paradigm. We developed four spatial normative frameworks of rsfMRI metrics (ALFF, fALFF, ReHo, and Hurst exponent), each capturing a distinct aspect of the BOLD signal, as a function of age and sex, accounting for MRI scan-related effects.

However, this work is broad but not comprehensive. Other rsfMRI metrics can provide additional features of the BOLD signal. Further inclusion of more metrics would enrich the multifaceted characterization of the BOLD signal. Moreover, this initial framework focused on a relatively narrow age window with an unbalanced sex ratio. Further work is needed to characterize the normative trajectories across a wider lifespan and more balanced sex ratios, which would allow further investigation of age-and sex-related effects on the BOLD signal. We believe these future directions are becoming increasingly feasible with the growing availability of rsfMRI databases. Future work should also investigate how different rsfMRI metrics, when used jointly within a normative framework, can provide complementary information on brain function.

Overall, this normative characterization of rsfMRI metrics in young population provides a reference across several features of the BOLD signal that indirectly reflect brain function. The development of these frameworks has the potential to improve the understanding of brain function while providing a reference to assess heterogeneity and individual deviations. Future work should apply these models to clinical cohorts and link individual deviations to clinically relevant outcomes to establish their importance to disease.

## Acknowledgements

This work was supported by the Natural Sciences and Engineering Research Council (NSERC) of Canada Discovery grant (grant number RGPIN-2023-05698) (to M. Noseworthy).

## Supporting information captions

**S1 Figure: Normative trajectories across age for each rsfMRI metric, across all ROIs**. All plots show five centile trajectories: 5^th^ and 95^th^ (dotted line, thin line width); 25^th^ and 75^th^ (dashed, medium line width), and 50^th^ (i.e., median; solid, thick line width). Sex-specific centile curves are displayed, with blue denoting males and orange denoting females. L: left; R: right.

**S1 Table**: **Summary of databases and their corresponding sites**. This table shows the sample size, sex distribution, MRI manufacturer, and rsfMRI sequence parameters, including TR, TE, number of time points, voxel size, number of slices, and eye condition. NA values indicate no information available. A mixed eyes condition indicates that the study has mixed instructions between eyes open and eyes closed. All studies were collected in 3T MRI systems.

**S2 Table: Summary of evaluation statistics for each rsfMRI metric normative model**. The reported statistics are (1) explained variance (EXPV); (2) Pearson’s correlation coefficient (RHO); (3) Standardized mean squared error (SMSE); (4) Shapiro-Wilk test; and (5) Mean standardized log loss (MSLL). The interpretation of these statistics is shown in **Table 1**.

